# Age-Related Differences in Ventral Striatal and Default Mode Network Function During Reciprocated Trust

**DOI:** 10.1101/2021.07.29.454071

**Authors:** Dominic S. Fareri, Katherine Hackett, Lindsey J. Tepfer, Victoria Kelly, Nicole Henninger, Crystal Reeck, Tania Giovannetti, David V. Smith

**Affiliations:** Gordon F. Derner School of Psychology, Adelphi University, Garden City, NY, USA; Department of Psychology and Neuroscience, Temple University, Philadelphia, PA, USA; Department of Psychological and Brain Sciences, Dartmouth College, Hanover, NH, USA; Lew Klein College of Media and Communication, Temple University, Philadelphia, PA, USA; Fox School of Business, Temple University, Philadelphia, PA, USA

**Keywords:** Trust, reciprocity, aging, ventral striatum, default mode network, connectivity

## Abstract

Social relationships change across the lifespan as social networks narrow and motivational priorities shift to the present. Interestingly, aging is also associated with changes in executive function, including decision-making abilities, but it remains unclear how age-related changes in both domains interact to impact financial decisions involving other people. To study this problem, we recruited 50 human participants (N_younger_ = 26, ages 18-34; N_older_ = 24, ages 63-80) to play an economic trust game as the investor with three partners (friend, stranger, and computer) who played the role of investee. Investors underwent functional magnetic resonance imaging (fMRI) during the trust game while investees were seated outside of the scanner. Building on our previous work with younger adults showing both enhanced striatal responses and altered default-mode network (DMN) connectivity as a function of social closeness during reciprocated trust, we predicted that these relations would exhibit age-related differences. We found that striatal responses to reciprocated trust from friends relative to strangers and computers were blunted in older adults relative to younger adults, thus supporting our primary pre-registered hypothesis regarding social closeness. We also found that older adults exhibited enhanced DMN connectivity with the temporoparietal junction (TPJ) during reciprocated trust from friends compared to computers while younger adults exhibited the opposite pattern. Taken together, these results advance our understanding of age-related differences in sensitivity to social closeness in the context of trusting others.

## Introduction

The transition to older adulthood is dynamic, characterized by shrinking social networks, prioritization of social relationships and socially-centered goals, and an emphasis on positive relative to negative social experiences (Carstensen, 1992, 1995; Charles, 2010). At the same time, there may also be benefits to interacting with less close individuals in older adulthood, since we may lose members of our inner social networks as we age. For example, having larger numbers of weaker social ties as we enter older adulthood is more strongly predictive of lower levels of depression and higher levels of positive affect 10-15 years later than the number of close relationships one has (Huxhold et al., 2020). These age-related socioemotional changes are coupled with changes in neural networks supporting executive and social function (Andrews-Hanna et al., 2007; Devitt and Schacter, 2020; Hughes et al., 2020; Laurita et al., 2020; Persson et al., 2006; Spreng et al., 2020). Taken together, these patterns have implications for older adults’ abilities to successfully engage in social interactions.

While an emerging body of literature has highlighted changes in decision-making and executive function in older relative to younger adults (Lighthall et al., 2018; Seaman et al., 2016; Burr et al., 2021) we know surprisingly little about how this translates to *social* decisions (i.e., trusting others). Extant findings suggest that older adults show increased rates of decisions to trust others, relative to younger adults (Bailey et al., 2015), and are less sensitive to having their decisions to trust be shaped by concerns of reputation. Older adults are also more likely to perceive others as more trustworthy relative to younger adults and are less likely to update their impressions of others based on subtle signs of dishonesty during social interactions (Bailey et al., 2016; 2019), potentially relying more heavily on initial facial appraisals (Suzuki, 2018). Yet, work examining whether age-related differences emerge in decisions to trust close others, relative to strangers, and in the processing of reciprocity and betrayal is understudied and has implications for understanding both how we integrate social- and value-related information with age.

Building and maintaining social relationships across the lifespan requires the ability to appraise someone as (un)trustworthy and effectively interpret their behavior within social interactions (Fareri, 2019). Research in young adults indicates that learning to trust draws on initial impressions of others that are dynamically updated with experienced patterns of reciprocity, a social reward signal (Chang et al., 2010); this process consistently recruits reward-related neural circuits (e.g., ventral striatum, medial prefrontal cortex; Bellucci et al., 2016; Fareri et al., 2012; Fouragnan et al., 2013). Moreover, our prior work (Fareri et al., 2015) demonstrates that these circuits differentially encode the value of reciprocity as a function of social closeness: reciprocity from friends relative to strangers elicits enhanced activation of the ventral striatum. This increased value signaling may in turn reinforce already close bonds, and in fact, decisions to trust in younger adults are strongly related to beliefs about safety (Chen et al., 2021). Yet, how social outcome processing shifts as we transition to older adulthood remains an outstanding question.

Critically, trust-based interactions are also predicated on an ability to create and adapt models of others and their intentions. Such theory-of-mind processes engage a network of brain regions including the temporoparietal junction (TPJ), posterior cingulate cortex (PCC), and medial prefrontal cortex (mPFC). Together, these regions comprise the “social brain,” which shares substantial overlap with the default-mode network (DMN; (Mars et al., 2012)). The DMN shows enhanced reactivity to social relative to non-social outcomes (Fareri et al., 2020) and may serve to prime us for engagement with the social world (Meyer, 2019). One possibility is that functional alterations within the DMN or between the DMN and reward-related networks may underlie the increased positivity and focus on beneficial social engagement in older adults, in turn contributing to an increased risk for exploitation (Castle et al., 2012; Harlé and Sanfey, 2012; Spreng et al., 2017).

Here, we sought to systematically investigate age-related differences in the effects of social closeness on trust behavior and the neural representation of reciprocity. In a pre-registered study, we implemented a variant of a trust game task used in previous work from our group in younger adults (Fareri et al., 2015): participants played with a computer, a stranger, and a close friend who accompanied them to the experiment. Based on findings from this study, we expected that participants overall would invest more with close friends relative to strangers and computers (pre-registered hypothesis 1.1); we further expected this differential pattern to be blunted in older (ages 63-80) relative to younger (ages 18-34) adults, given increased rates of trust behavior in older adults overall (e.g., Bailey et al., 2015). In line with our behavioral predictions, we additionally expected blunted striatal responses to reciprocity from friends relative to strangers in older adults (pre-registered hypothesis 2.1). We also hypothesized that age-related differences in striatal responses to reciprocity would be tied to DMN-striatal connectivity (pre-registered hypothesis 2.1), which in turn would mediate expected age-related differences in trust behavior (pre-registered hypothesis 2.2); we expected this mediation to be moderated by self-reported social closeness with friends relative to strangers in the trust game (pre-registered hypothesis 2.3).

## Methods

### Participants

Fifty participants (26 young adults, ages 18-34; 24 older adults, ages 63-80) were recruited to participate in this study. This sample size was pre-registered (https://aspredicted.org/MVZ_ODI), determined a priori before data collection, and limited largely by available funding for data collection for this project. We acknowledge that the relatively small sample size limits our ability to draw strong inferences, especially in analyses relating brain activation and self-reports of social closeness. Young adult participants were recruited primarily through the Temple University Psychology Department participant pool and received course credit (along with a task bonus in gift cards) for participation. Older adult participants were recruited using a range of efforts—reaching out to local community and senior centers, newspaper advertisements, and local flyers—and were compensated with Amazon gift cards for their participation ($25 per hour of participation for MRI participants, $15 per hour for both their friends and recruited confederates to act as strangers; bonus payment for MRI participants and their friends varied across individuals based on randomly chosen outcomes paid out at the end of the experimental session). All participants were screened before data collection to rule out current major psychiatric or neurologic illness, as well as MRI contraindications. Older adults were screened to rule out dementia using the Telephone Interview for Cognitive Status, with a score under 30 meeting exclusion criteria (Brandt et al., 1988). All participants included in analyses had at least 2 usable runs of the task. These exclusions left a final sample of 48 total participants, with 26 younger adults (mean age: 23.2 years, *SD:* 4.07 years; 35% male) and 22 older adults (mean age: 69.3 years; SD: 4.38 years; 50% male). All participants gave written informed consent as part of a protocol approved by the Institutional Review Board of Temple University.

### Procedures

Participants completed two appointments. During the first appointment, participants underwent a mock MRI scan to help control for motion and acclimate to the scanner. They also completed a brief neuropsychological test battery including measures of estimated premorbid intelligence, specific cognitive domains (e.g., attention, executive functioning, episodic memory, language), and self-reported everyday functioning (older adults only). Participants completed a second appointment with a self-selected friend of the same identified sex and age group (within 5-10 years, who was not a family member or spouse). During the second appointment, participants completed a 45-minute survey that included measures of mood, media usage, and emotion regulation (not presented here) as well as questions about perceived social closeness with their friend and the stranger (Inclusion of Other in Self Scale (IOS); Aron et al., 1992), and basic demographics.

### Experimental Paradigm

After completing the survey, participants and their friends were introduced to a sex- and age-matched (within 10 years) stranger (confederate) and were then trained on the Trust Game task (adapted from Fareri et al., 2015)). MRI participants were told that they would be playing a game called the investment game in real time with their friend, the stranger and a computer partner. On a given trial of the task, the MRI participant would play with one of their three partners as indicated by a photo and name presented on the screen. Participants were instructed that they would start each trial with $8 and that they would have a choice between sending (investing) different proportions of that $8 to their partner on a given trial. The amounts that could be sent varied on a trial to trial basis, ranging from $0-$8. Participants would have up to 3 seconds to indicate via a button press on an MRI compatible response box which of the two investment options they preferred. Participants were instructed that whatever amount they chose to invest would be multiplied by a factor of 3 (i.e., an investment of $6 would become $18 for the partner), and that their partner could decide to split the multiplied amount evenly with them (reciprocate) or keep it all for themselves (defect). Upon entering their response, participants would see a screen that said ‘waiting’, (1.5s) during which time they believed that their decision was being presented to their partner in another room in the research suite. After the waiting screen, a variable ISI was presented (mean = 1.42s), and participants were then notified (2s) whether their partner decided to split that amount evenly with them (reciprocate) or keep (defect) all of the money. Unbeknownst to participants, all outcomes were predetermined, and all partners were preprogrammed to reciprocate 50% of the time as per our previous work (Fareri et al., 2015).

After task training, the MRI participant, the stranger and close friend were split up into different rooms before beginning the actual experimental session. Participants underwent a 90-minute MRI scan that included up to 5 runs of the trust game task, as well as two additional, but separate tasks (not discussed here). Each run of the task consisted of 36 trials, with 12 trials per partner. See Figure 1A and our previous work (Fareri et al., 2015) for task details. We note that after completing the trust game task, participants completed two separate, additional fMRI tasks aimed at investigating age-related differences in shared reward processing and bargaining behavior. However, the trust game task was the primary focus of the experimental session, and was always completed first.

**Figure 1:**
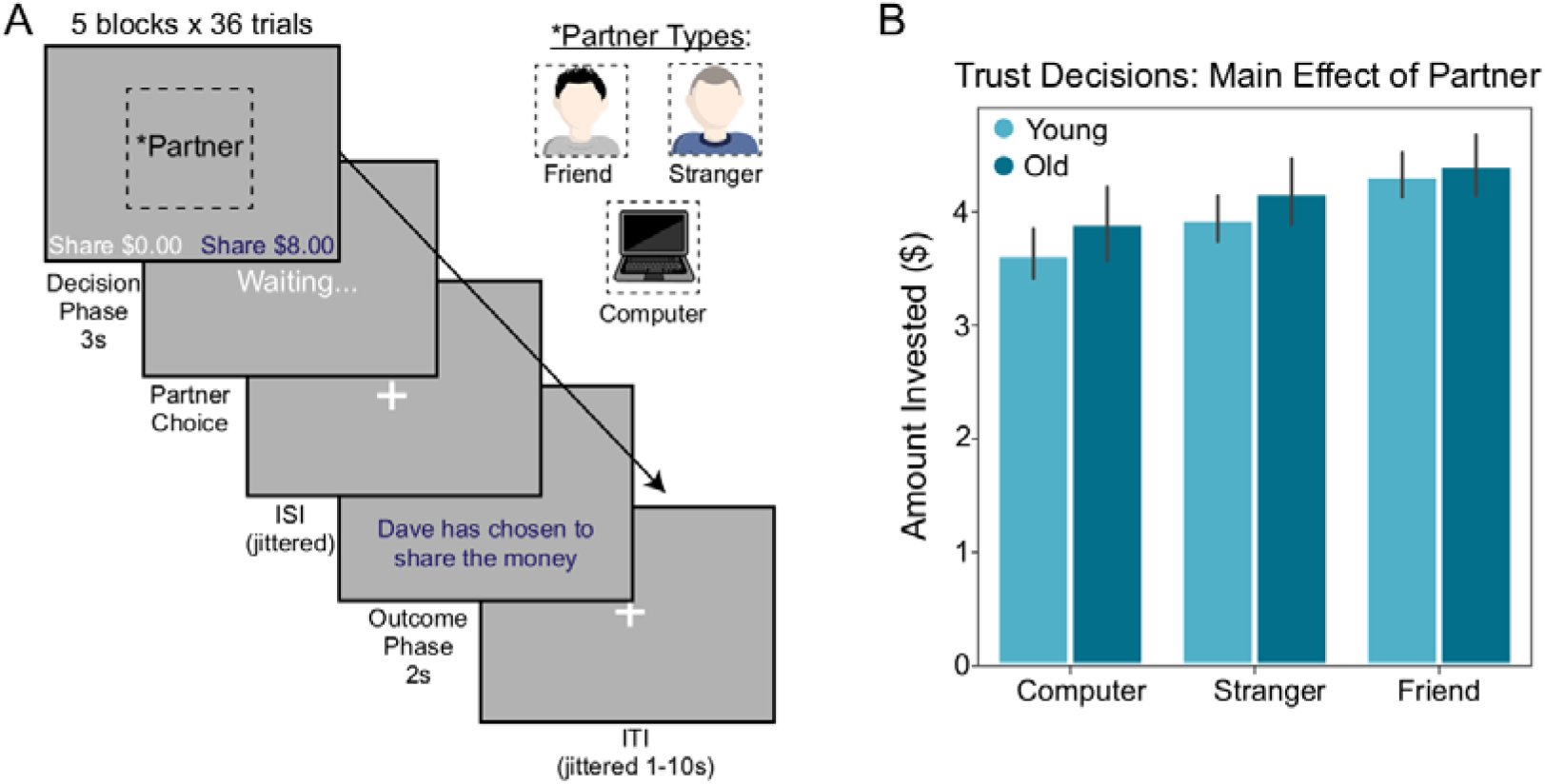
Task and Behavior. (A) We used an economic trust game to measure investments made to three different types of partners. Participants played the role of the investor with three distinct investees of varying social closeness (friend, stranger, and computer as a nonsocial control). On each trial of the trust task, participants were first shown an image of their partner with a neutral expression (note that images shown here are for illustrative purposes only; the experiment utilized real pictures of the partners) and were given a choice to invest a greater or lesser amount of money with their partner. Money invested into the partner triples. When money was invested into the investee, the participant was then shown a screen indicating that the investee is contemplating the investment (“Waiting…”). After a brief jittered ISI, the participant is shown the outcome of the investee’s decision (i.e., reciprocate and share the money evenly, or defect and take all of the money). (B) A 3 (partner) x 2 (age group) repeated measures ANOVA revealed that both older and younger participants differentially invested money as a function of partner.

### Behavioral Analyses

To ensure that we effectively manipulated social closeness, we conducted a 2 (partner) x 2 (age group) repeated measures ANOVA on participants’ IOS ratings of their friends and strangers. We examined participants’ decisions to trust (i.e., invest) as a function of partner and age by conducting a 3 (Partner) x 2 (Age Group) repeated measures ANOVA (see ‘Deviations from Pre-Registration’ section in Supplementary Methods). We also conducted this analysis controlling for individual differences in reaction time by z-scoring participants’ mean reaction time during the decision phase and including these values as a covariate in an ANCOVA. This was done to ensure any potential differences in trust decisions were not accounted for by differences in reaction time during decision-making. We similarly performed an exploratory 3 (Partner) x 2 (Age Group) repeated measures ANOVA solely on participants’ reaction time during the decision phase. While there are well-documented differences in reaction time as a function of age (Salthouse, 2000; Finkel, 2007), we were interested in exploring if and potentially how such differences might change as a function of social partner in this task. We also performed additional exploratory analyses assessing differences in choice behavior and reaction time as a function of whether participants experienced reciprocity or a violation of trust on the previous trial as a means to probe the influence of experienced outcomes on trust behavior.

### Neuroimaging Data Acquisition

Neuroimaging data were collected at the Temple University Brain Research and Imaging Center (TUBRIC) using a 3.0 Tesla Siemens Prisma scanner equipped with a 20-channel phased array head coil. Functional images sensitive to blood-oxygenation-level-dependent (BOLD) contrast were acquired using a single-shot T2*-weighted echo-planar imaging sequence with slices roughly parallel to the axial plane collected in descending order [repetition time (TR): 2.02 s; echo time (TE): 23 ms; matrix 74 x 74; voxel size: 2.97 x 2.97 x 2.80 mm; 36 slices (15% gap); flip angle: 76°]. To facilitate co-registration and normalization of functional data, we also collected high-resolution T1-weighted structural scans (TR: 2.4 s; TE: 2.2 ms; matrix 192 x 192; voxel size: 1.0 mm^3^; 192 slices; flip angle: 8°) and B_0_ field maps (TR: 645 ms; TE_1_: 4.92 ms; TE_2_: 7.38 ms; matrix 74 x 74; voxel size: 2.97 x 2.97 x 2.80 mm; 36 slices, with 15% gap; flip angle: 60°). In addition, we also collected T2-weighted structural images (TR: 3.2 s; TE: 567 ms; matrix 192 x 192; voxel size: 1.0 mm^3^; 192 slices; flip angle: 120°); these images are included with our data on OpenNeuro.org, but we did not use them in our preprocessing or analyses.

### Preprocessing of Neuroimaging Data

Neuroimaging data were converted to the Brain Imaging Data Structure (BIDS) using HeuDiConv version 0.5.4 (Halchenko et al., 2019). Results included in this manuscript come from preprocessing performed using *fMRIPrep* 20.1.0 (Esteban et al., 2018b, 2018a), which is based on *Nipype* 1.4.2 (Gorgolewski et al., 2011, 2018). The details described below are adapted from the *fMRIPrep* preprocessing details; extraneous details were omitted for clarity.

#### Anatomical data preprocessing

The T1-weighted image was corrected for intensity non-uniformity (INU) with N4BiasFieldCorrection (Tustison et al., 2010), distributed with ANTs 2.2.0 (Avants et al., 2008), and used as T1w-reference throughout the workflow. The T1w-reference was then skull-stripped with a *Nipype* implementation of the antsBrainExtraction.sh workflow (from ANTs), using OASIS30ANTs as target template. Brain tissue segmentation of cerebrospinal fluid (CSF), white-matter (WM) and gray-matter (GM) was performed on the brain-extracted T1w using FAST (FSL 5.0.9, (Zhang et al., 2001)). Volume-based spatial normalization to MNI152NLin2009cAsym standard space was performed through nonlinear registration with antsRegistration (ANTs 2.2.0), using brain-extracted versions of both T1w-reference and the T1w template. To this end, the *ICBM 152 Nonlinear Asymmetrical template version 2009c* (Fonov et al., 2009) template was selected for spatial normalization.

#### Functional data preprocessing

For each of the BOLD runs contained per subject, the following preprocessing steps were performed. First, a reference volume and its skull-stripped version were generated using a custom methodology of *fMRIPrep* (Esteban et al., 2018b). Head-motion parameters with respect to the BOLD reference (transformation matrices, and six corresponding rotation and translation parameters) are estimated before any spatiotemporal filtering using mcflirt (FSL 5.0.9, (Jenkinson et al., 2002)). BOLD runs were slice-time corrected using 3dTshift from AFNI 20160207 (Cox and Hyde, 1997). A B0-nonuniformity map (or *fieldmap*) was estimated based on a phase-difference map calculated with a dual-echo GRE (gradient-recall echo) sequence, processed with a custom workflow of *SDCFlows* inspired by the <u>epidewarp.fsl script</u> and further improvements in HCP Pipelines (Glasser et al., 2013). The *fieldmap* was then co-registered to the target EPI (echo-planar imaging) reference run and converted to a displacements field map (amenable to registration tools such as ANTs) with FSL’s fugue and other *SDCflows* tools. Based on the estimated susceptibility distortion, a corrected EPI (echo-planar imaging) reference was calculated for a more accurate co-registration with the anatomical reference. The BOLD reference was then co-registered to the T1w-reference using FLIRT (FSL 5.0.9, (Jenkinson and Smith, 2001)) with the boundary-based registration (Greve and Fischl, 2009) cost-function. Co-registration was configured with nine degrees of freedom to account for distortions remaining in the BOLD reference. The BOLD time-series (including slice-timing correction when applied) were resampled onto their original, native space by applying a single, composite transform to correct for head-motion and susceptibility distortions. These resampled BOLD time-series will be referred to as *preprocessed BOLD in original space*, or just *preprocessed BOLD*. The BOLD time-series were resampled into standard space, generating a *preprocessed BOLD run in MNI152NLin2009cAsym space*.

Additionally, a set of physiological regressors were extracted to allow for component-based noise correction (CompCor, (Behzadi et al., 2007)). Principal components are estimated after high-pass filtering the preprocessed BOLD time-series (using a discrete cosine filter with 128s cut-off) for the two CompCor variants: temporal (tCompCor) and anatomical (aCompCor). tCompCor components are then calculated from the top 5% variable voxels within a mask covering the subcortical regions. This subcortical mask is obtained by heavily eroding the brain mask, which ensures it does not include cortical GM regions. For aCompCor, components are calculated within the intersection of the aforementioned mask and the union of CSF and WM masks calculated in T1w space, after their projection to the native space of each functional run (using the inverse BOLD-to-T1w transformation). Components are also calculated separately within the WM and CSF masks. For each CompCor decomposition, the k components with the largest singular values are retained, such that the retained components’ time series are sufficient to explain 50 percent of variance across the nuisance mask (CSF, WM, combined, or temporal). The remaining components are dropped from consideration. As an additional confound, we also estimated framewise displacement (FD). FD was computed using the relative root mean square displacement between affines (Jenkinson et al., 2002).

All resamplings can be performed with *a single interpolation step* by composing all the pertinent transformations (i.e., head-motion transform matrices, susceptibility distortion correction when available, and co-registrations to anatomical and output spaces). Gridded (volumetric) resamplings were performed using antsApplyTransforms (ANTs), configured with Lanczos interpolation to minimize the smoothing effects of other kernels (Lanczos, 1964).

### Neuroimaging Analyses

Neuroimaging analyses used FSL version 6.03 (Smith et al., 2004; Jenkinson et al., 2012). We specifically focused on two types of models (activation and connectivity) to quantify how reciprocated trust and social closeness were associated with BOLD responses. Both types of models were based on a general linear model with local autocorrelation (Woolrich et al., 2001). Our first model focused on the brain activation evoked during the trust task and used a total of nine regressors of interest. We used three regressors to model the decision phase (duration = response time) associated with each of the partners (computer, stranger, friend). To model brain activation associated with outcomes (reciprocate and defect) by partner, we used six additional regressors (duration = 1 second). Each task-related regressor was convolved with the canonical hemodynamic response function. We also conducted offline analyses examining the relation between reward-related BOLD responses and self-reported social closeness (see Deviations from Pre-registration section, Supplementary Methods).

Our second type of model focused on the task-dependent connectivity associated with the trust task. To estimate these changes in connectivity, we used psychophysiological interaction (PPI) analysis (Friston et al., 1997; O’Reilly et al., 2012). Regions that exhibit a significant PPI effect can be interpreted as showing a context-specific modulation of effective connectivity, though the directionality of this modulation remains ambiguous without additional analyses (Smith et al., 2016; Friston et al., 2003). Notably, recent meta-analytic work has shown that PPI reveals consistent and specific patterns of connectivity across multiple seed regions and psychological contexts (Smith et al., 2016; Smith and Delgado, 2017). We first estimated a network PPI model that focused on task-dependent changes in connectivity with the DMN (Utevsky et al., 2017; Fareri et al., 2020). The DMN and nine additional networks, including the executive control network (ECN) were based on prior work (Smith et al., 2009). Network time courses were extracted with a spatial regression component of the dual regression approach (Filippini et al., 2009; Nickerson et al., 2017) and entered into a model with the nine task regressors from the activation model described above. PPI regressors were formed by multiplying each of the nine task regressors by the DMN regressor, yielding a total of 28 regressors.

We also conducted exploratory seed-based analyses in regions that extant research implicates in reward and social processes (Fareri et al., 2012; 2015; 2020; Utevsky et al., 2017; Smith et al., 2016, 2010; Chib et al., 2018; Tomova et al., 2020), including the ventral striatum, vmPFC, FFA, TPJ and PCC (see Results). All seed regions were functionally defined from our wholebrain analyses, and can be found on Neurovault (https://neurovault.org/collections/10477). For each participant in these analyses, the average time course of a seed was extracted and entered as a regressor into a model with the nine task regressors from the activation model described above. PPI regressors were formed by multiplying each of the nine task regressors by the seed regressor, yielding a total of 19 regressors in each seed-based PPI model.

Both activation and connectivity models included a common set of confound regressors. We first modeled out missed responses by including an additional task-related regressor to account for the full duration of the choice screen. We also included additional regressors for the six motion parameters (rotations and translations), the first six aCompCor components explaining the most variance, non-steady state volumes, and the framewise displacement (FD) across time. Finally, high-pass filtering (128s cut-off) was achieved using a set of discrete cosine basis functions.

We combined data across runs, for each participant, using a fixed-effects model. Group-level analysis was carried out using FLAME (FMRIB’s Local Analysis of Mixed Effects) Stage 1 and Stage 2 (Beckmann et al., 2003; Woolrich et al., 2004). Our group-level model focused on comparisons between older and younger groups; these comparisons included covariates to account for gender, temporal signal to noise ratio (tSNR), mean framewise displacement, and mean response time. All z-statistic images were thresholded and corrected for multiple comparisons using an initial cluster-forming threshold of z□>□3.1 followed by a whole-brain corrected cluster-extent threshold of p□<□ü.05, as determined by Gaussian Random Field Theory (Worsley, 2001).

#### Data and code availability

Data can be found on https://openneuro.org/datasets/ds003745. Analysis code can be found on https://github.com/DVS-Lab/srndna-trustgame. Thresholded and unthresholded statistical maps are located on https://identifiers.org/neurovault.collection:10447.

## Results

Below we report results from behavioral analyses and both task-based neural activation and connectivity analyses. We begin by presenting results of manipulation checks regarding selfreported social closeness, followed by results of pre-registered and exploratory behavioral analyses targeting age-related differences in trust behavior. We then present results of preregistered and exploratory task-based fMRI activation analyses targeting age differences in processing of trust game outcomes. Last, we present results of pre-registered and exploratory task-based connectivity (PPI) analyses examining age-related differences in communication within reward-related and social neural systems.

### Self-reported social closeness is greater for friends relative to strangers

As a manipulation check, we investigated whether participants exhibited differences in selfreported social closeness to friends and strangers. A 2×2 repeated-measures ANOVA on IOS scores revealed a significant main effect of partner (*F_(1,41)_* = 70.40, *p* < .001), such that participants reported feeling closer to friends (M = 4.50, SD = 1.70) relative to strangers (M = 2.14, SD = 1.52). Contrary to our hypotheses, we observed neither a significant interaction of partner and age group (*F_(1,41)_* = 0.10, *p* = 0.76), nor a significant main effect of age group (*F_(1,41)_* = 2.53, *p* = .12). We note that due to technical error, a number of participants (n = 5) lacked closeness ratings for the stranger and were unable to be included in this analysis.

### Social closeness shapes trust decisions

Our first goal was to examine differences in trust behavior as a function of age and partner. To test pre-registered hypothesis 1.1, we conducted a 3 (partner) x 2 (age group) repeated measures ANOVA on the average amount of money invested during the trust game. Controlling for individual differences in overall reaction time, we observed partial support for pre-registered hypothesis 1.1: a significant main effect of partner on amount of money invested emerged (*F_(2,90)_* = 18.49, *p* < .001, □^2^_p_ = .291; see Figure 1B), such that participants sent more money to their friends relative to both strangers (*t_(88)_* = 2.86, *p* < .01) and computers (*t_(88)_* = 5.14, *p* < .001). This result replicates prior work from our group (Fareri et al., 2015). However, inconsistent with hypothesis 1.1, we did not observe an interaction between partner and age (*F_(2,90)_* = 2.16, *p* = 0.12, □^2^_p_ = .046), nor did we observe a significant difference between older and younger adults in amount of money invested overall (*F_(1,45)_* = 2.77, *p* = .10, □^2^_p_ = .058). An additional exploratory ANOVA on choice behavior following reciprocity and violations of trust revealed that participants overall invested more with friends relative to other partners regardless of previous outcome (*F_(2,92)_* = 7.38, *p* <.001, □^2^_p_ = .138) and invested more after reciprocity relative to violations of trust (*F_(1,46)_* = 10.74, *p*<.002, □^2^_p_ = .189). We note that no significant interactions emerged between either factor and age, or between partner, choice and age.

We also conducted an exploratory 3 (partner) x 2 (age group) repeated measures ANOVA on reaction time. This analysis revealed a significant effect of age (*F_(1,46)_* = 7.60, *p* < .01, □^2^_p_ =.142), with older adults exhibiting significantly longer reaction times when making their decisions, but no significant effect of partner (*F_(2,92)_* = 2.75, *p* = .069, □^2^_p_ =.056).

An additional exploratory ANOVA on reaction time as a function of outcomes on previous trials revealed that participants responded more quickly on trials with friends relative to other partners (*F_(2,92)_* = 3.08, *p* = .05, □^2^_p_ = .063) and more quickly after experiencing reciprocity (*F_(1,46)_* = 15.58, *p* <.001, □^2^_p_ = .253). We interestingly observed a significant three-way interaction between partner, choice and age on reaction time (*F_(2,92)_* = 3.16, *p* < .05, □^2^_p_ = .064), whereby younger adults show similarly quick reaction times after either reciprocity or violations of trust with all partners, whereas older adults show longer reaction times after experiencing violations of trust from computer partners, but similar reaction times regardless of previous outcome with social partners.

Given that there are well-documented age-related differences in processing speed and reaction times (Salthouse, 2000; Finkel, 2007), all of our analyses sought to account for these effects by including additional covariates related to response time. Nevertheless, we also note that global age-related differences in RT could be tied to changes in behavior. In order to account for this possibility, we re-ran exploratory analyses focusing on post-outcome shifts in behavior including participants’ average RT (collapsed across partner conditions) as a covariate. Results focusing on investment rates as a function of previous outcomes held when including this additional covariate (main effect of partner: F_(2,86)_ = 7.26, p < .001, □^2^p = .144; main effect of previous outcome: F_(2,86)_ = 9.73, p < .005, □^2^p = .184). Results focusing on reaction time as a function of previous outcomes partially held (main effect of partner: F_(2,86)_ = 3.97, p < .03, □^2^p = .085; main effect of previous outcome: F_(2,86)_ = 12.72, p < .001, □^2^p = .228), but we note that the partner x previous outcome x age interaction was no longer significant (F_(1,43)_ = 2.13, p = .126, □^2^p = .047).

#### Blunted striatal responses to reciprocity from friends in older adults

Based on our expected behavioral results and our prior work, we were also interested in characterizing differences in task-based neural activation and connectivity specifically during experienced outcomes (i.e., reciprocity vs. defection of trust) in the trust game. We first hypothesized that participants would exhibit enhanced reward-related responses within the striatum when experiencing reciprocity from a close friend relative to other partners, and that this effect would be stronger in younger adults and associated with relationship closeness (pre-registered hypothesis 2.1). To test this hypothesis, we conducted a confirmatory whole-brain contrast of reciprocate > defect. This analysis revealed robust activation of the bilateral ventral striatum (Rt. ventral striatum: x, y, z = 7.5, 13.2, −4.4; Lt. ventral striatum: x, y, z = −10.3, 16.2, −1.2; see Fig. 2A), among other regions, including vmPFC, mPFC, PCC and occipital cortex (see Table 1 and https://identifiers.org/neurovault.image:512088 for more information).

**Figure 2:**
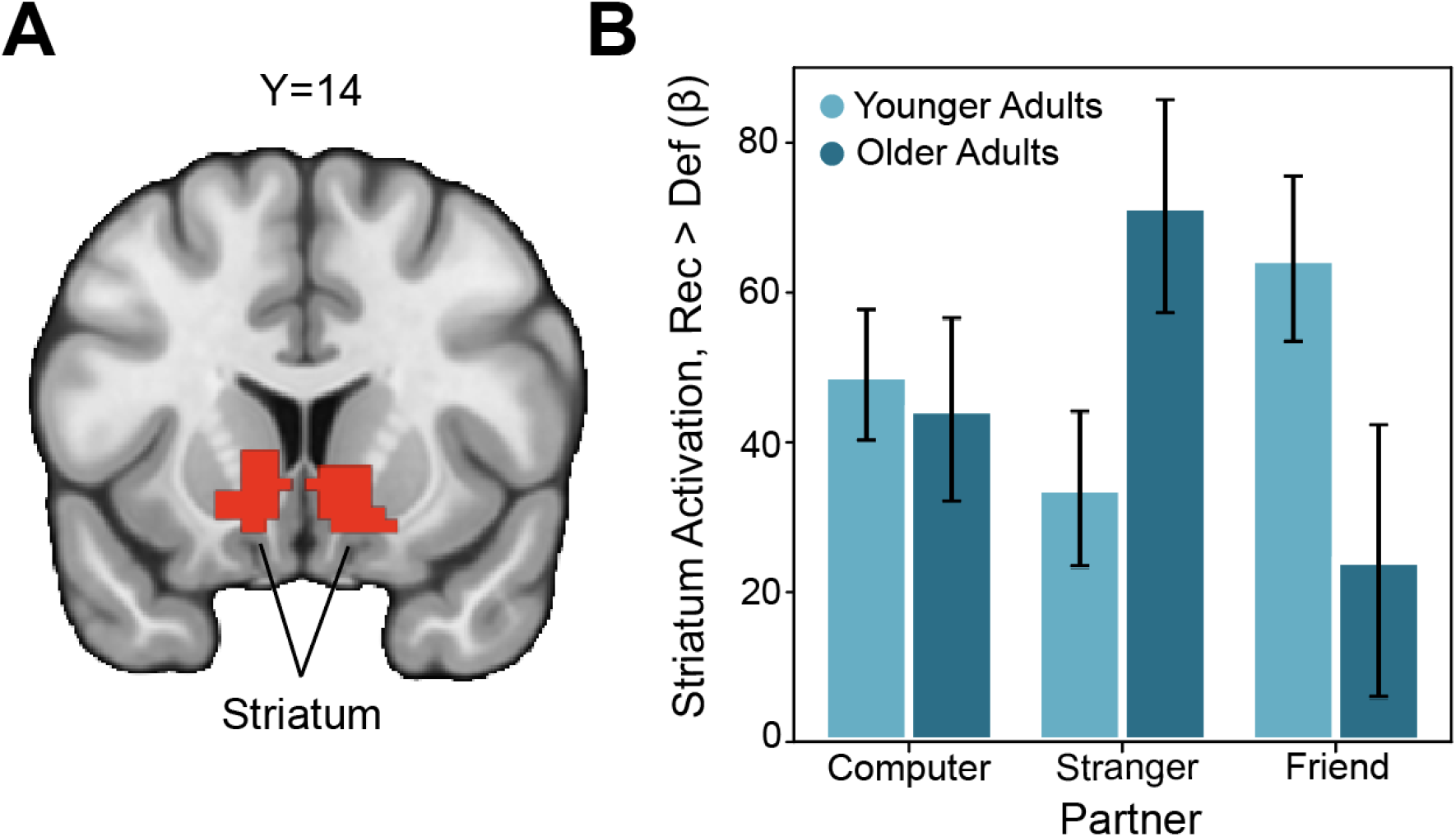
Blunted Effects of Social Closeness on Striatal Social Reward Responses in Older Adults. (A) We identified voxels in the striatum whose activation increased for reciprocate outcomes relative to defect outcomes, irrespective of age. The overlaying Z statistic image was thresholded parametrically (Gaussian Random Field Theory) using GRF-theory-based maximum height thresholding (i.e., voxel level threshold) with a (corrected) significance threshold of P=0.05 (Thresholded: https://identifiers.org/neurovault.image:512088; Unthresholded: https://identifiers.org/neurovault.image:512063). (B) We interrogated this striatal region further to probe for group differences in the effects of partner on reward processing. Although younger adults exhibited an enhanced striatal response to the contrast of reciprocate > defect when the partner was a friend relative to a stranger, older adults exhibited the opposite effect, suggesting that the effects of social closeness on reward related neural responses are blunted in older adults.

We next conducted a 2 (age group) x 3 (partner) repeated-measures ANOVA on extracted parameter estimates from the bilateral ventral striatum to investigate age differences in processing of trust game outcomes. We observed a significant partner x group interaction (F(2,92)= 5.94, p = .004, partial eta^2^ = .114; see Figure 2B). Consistent with patterns demonstrated in prior work from our group (Fareri et al., 2015), young adults demonstrated increased striatal activation when experiencing reciprocity (relative to defection) from friends relative to strangers, though we note that this was not significant (t(46) = 1.89, p = .065). Interestingly, older adults exhibited diminished striatal activation when experiencing reciprocate relative to defect outcomes from friends as compared to strangers (t(46) = −2.68, p = .01). Rerunning these analyses including tSNR, mean framewise displacement (FD) and sex as covariates eliminated the partner x group interaction effect (*F_(2,92)_* = 2.30, *p* = .107, □^2^_p_ = .051).

Last, we examined whether differences in self-reported social closeness with friends and strangers was associated with differences in the striatal response to trust game outcomes with friends and strangers. A linear regression regressing the difference in striatal BOLD for reciprocate > defect for friends > strangers on the difference in reported social closeness with those same partners did not reveal a significant effect (b = 13.00, SE = 7.86, *t* = 1.66, *p* = .104), though we note that the pattern was in the hypothesized direction of a positive relationship between self-reported social closeness and the striatal BOLD response to reciprocity from friends relative to strangers. Taken together, the results regarding striatal function provide partial support for pre-registered hypothesis 2.1.

#### Altered cortical responses to violations of trust in older adults

We also conducted an additional exploratory whole-brain interaction contrast to highlight regions demonstrating age-related differences in responses to trust game outcomes. This analysis revealed significant clusters of activation in the anterior insula, posterior cingulate and both dorsal and ventral medial prefrontal cortex (see Table 1). Extracting parameter estimates from the insula (x, y, z = 37, 25, 8) revealed a reduced response to defect relative to reciprocate outcomes specifically in the stranger condition in older adults (Fig 3A, B). Older adults also demonstrated a reduced response to reciprocate relative to defect outcomes experienced with friends in the PCC (x, y, z = 4, −58, 28; Fig 3C, D).

**Figure 3:**
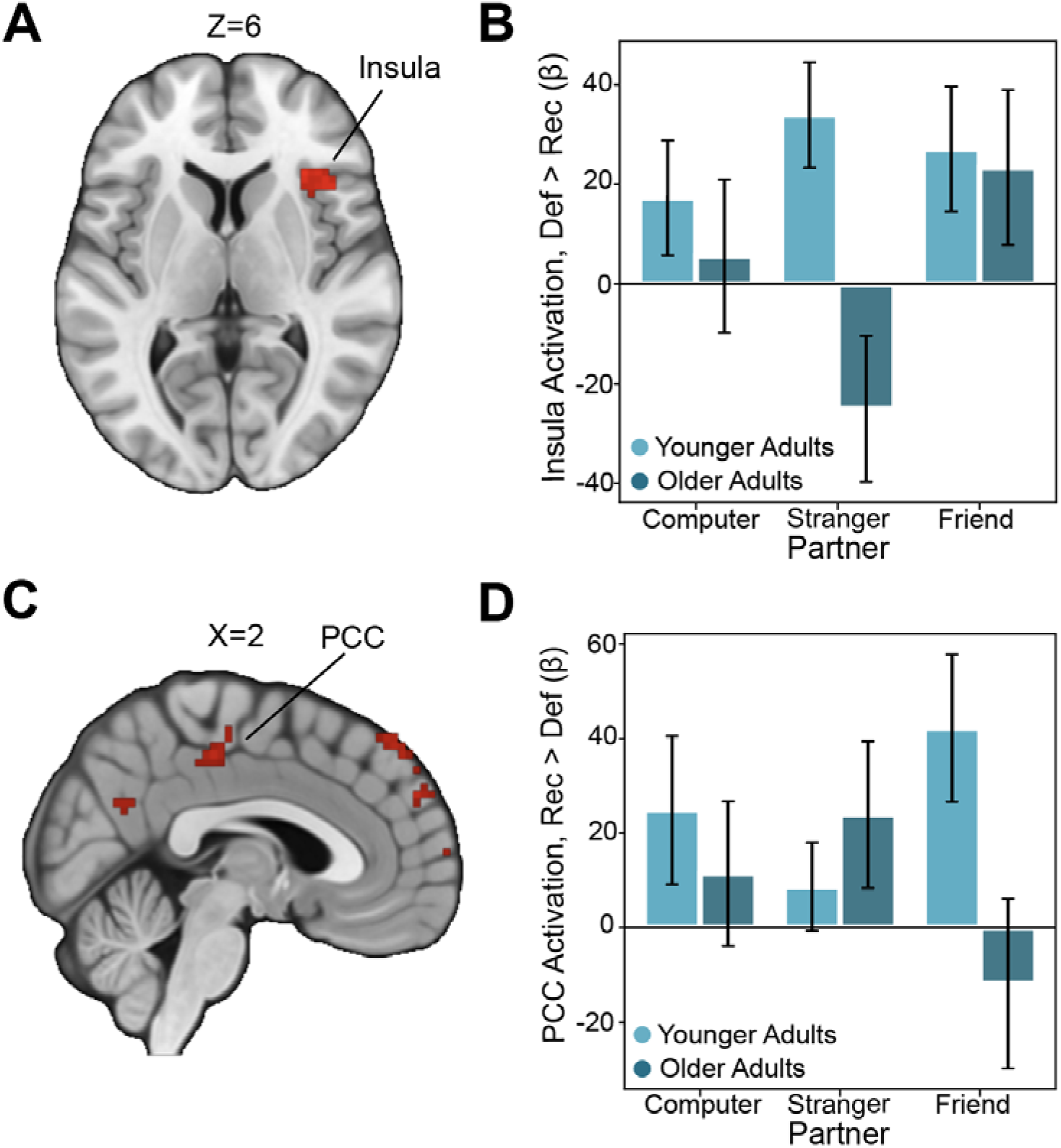
Age-related Differences in Neural Responses to Social Outcomes. (A) We found that the anterior insula exhibited a larger response to defect outcomes relative to reciprocate outcomes in younger adults (plotted here as defect > reciprocate), consistent with the idea that older adults are less sensitive to outcomes associated with losses (Thresholded: https://identifiers.org/neurovault.image:512048; Unthresholded: https://identifiers.org/neurovault.image:512076). (B) For descriptive purposes, we extracted beta values within this anterior insula region and confirmed that older adults showed a blunted response to defect outcomes. (C) In the opposite contrast (reciprocate > defect), we found several regions whose outcome-related activation was enhanced in younger adults (Thresholded: https://identifiers.org/neurovault.image:512037; Unthresholded: https://identifiers.org/neurovault.image:512065). (D) For descriptive purposes, we extracted the response within the posterior cingulate cortex (PCC) and confirmed that the response to reciprocation in this region was enhanced in younger adults. We note that Z statistic images were thresholded parametrically (Gaussian Random Field Theory) using clusters determined by Z>3.1 and a (corrected) cluster significance threshold of P=0.05.

#### Enhanced network connectivity during trust game outcomes with friends in older adults

Our task-based activation analyses revealed engagement of both reward related circuits and regions comprising the default mode network (e.g., PCC, mPFC) during experiences of reciprocity relative to violations of trust. As outlined in our pre-registration, we predicted that we would observe age-related differences in default mode connectivity, specifically with the striatum, when experiencing reciprocity relative to defection from different partners (H2.2). To test this prediction, we conducted a generalized network psychophysiological interaction (nPPI) analysis during the outcome phase of the task using the DMN as our seed network (cf. Fareri et al., 2020). A contrast of reciprocate > defect as a function of social closeness (friend > computer) and age (young > old) revealed enhanced connectivity between the DMN and TPJ (x, y, z = 64, −40, 24.5; see Fig. 4) when experiencing reciprocity from friends relative to the computer in older, compared to younger adults. This pattern of results may suggest that TPJ is more tightly integrated within the DMN in this particular task context. This analysis also revealed enhanced DMN coupling with the supplementary motor area and the occipital pole (see Table 1 and https://identifiers.org/neurovault.image:512032).

**Figure 4:**
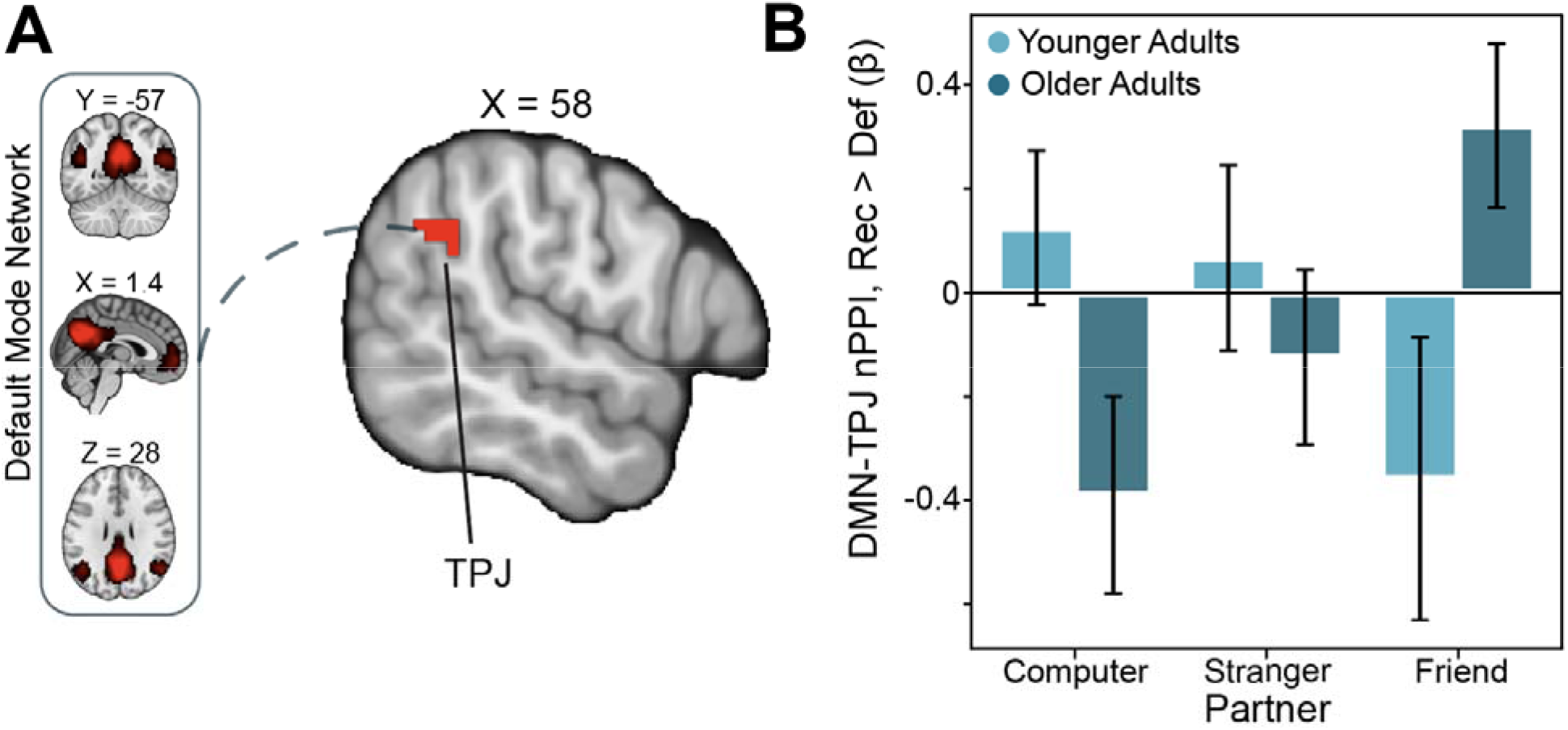
Older Adults Show Enhanced DMN-TPJ Connectivity as a Function of Social Closeness. (A) We used a network psychophysiological interaction (nPPI) analysis with the default mode network (DMN) as a seed. We examined connectivity with DMN as a function of age differences and social closeness (friend > computer) during the outcome phase (reciprocate > defect). This double subtraction analysis [(friend reciprocate > friend defect) > (computer reciprocate > computer defect)] indicated that DMN connectivity with the right temporal-parietal junction (TPJ) was enhanced in older adults (Thresholded: https://identifiers.org/neurovault.image:512032; Unthresholded: https://identifiers.org/neurovault.image:512035). (B) For illustrative purposes, we extracted the parameter estimates within this TPJ region. We note that Z statistic images were thresholded parametrically (Gaussian Random Field Theory) using clusters determined by Z>3.1 and a (corrected) cluster significance threshold of P=0.05.

Interestingly, we did not observe enhanced connectivity between the DMN and striatum as predicted in pre-registered hypothesis 2.2 and as such we did not pursue our final preregistered hypothesis (H2.3), which posited that DMN-striatal connectivity would mediate partner related effects in trust behavior. We did, however, conduct an additional exploratory analysis to probe whether the age-related differences in DMN-rTPJ connectivity during outcome processing was related to differences in trust behavior. We performed a linear regression, regressing DMN-TPJ connectivity estimates from a contrast of (Friend Reciprocate > Friend Defect) > (Computer Reciprocate > Computer Defect) on age (mean centered), the behavioral difference score (Friend investment - Computer investment, mean centered) and their interaction. Results of this analysis revealed a significant interaction of age and behavior on DMN-TPJ connectivity (b = −0.027, SE = 0.012, t = −2.22, p = 0.032; see Supplementary Figure 1), which suggest that the more younger adults invest with a friend relative to a computer, the more positive the connectivity is between DMN and TPJ when experiencing reciprocity (relative to defect outcomes) from friends relative to computers. Older adults demonstrate the opposite pattern; the age-related connectivity differences appear to be strongest when investing less with friends.

We also explored task-based connectivity of the Executive Control Network (ECN) during the processing of trust game outcomes. This exploratory analysis was motivated by previous work implicating the ECN in reward-processing and goal directed behavior more generally (Waltz et al., 2013; Fareri et al., 2020; Grill et al., 2021). We conducted a network PPI contrast of reciprocate > defect for friends > strangers as a function of age group. Here, we found that relative to younger adults, older adults demonstrated increased connectivity between the ECN and a region encompassing mid/posterior insula (x, y, z = 34, 4, −4; see Table 1) when experiencing defect outcomes with one’s friend, but not with the stranger or computer.

In a control analysis, we examined whether these network connectivity effects were due to the template networks being derived from a younger and independent sample (Smith et al., 2009). We therefore used independent component analyses to identify networks in our sample (see Supplementary Methods). Results from both control analyses (DMN-TPJ, ECN-insula) partially replicated the patterns observed in our whole brain results (see Supplementary Results).

#### Reduced vmPFC-hippocampus connectivity during reciprocity in older adults

Finally, we also conducted additional exploratory seed-based PPI analyses investigating group differences in responses to trust game outcomes as a function of social closeness using the following seeds: FFA, vmPFC and ventral striatum, TPJ and PCC. Finally, we also conducted additional exploratory seed-based PPI analyses using the following seed regions defined from contrasts of Reciprocate outcomes > Defect outcomes (vmPFC, ventral striatum) and Friend + Stranger decisions > Computer decisions (FFA, TPJ, PCC). We note that extant literature also implicates these regions in reward-related and social processes, respectively. We hypothesized that they may also exhibit age-related differences in their connectivity with other regions. Robust group differences emerged (see the NeuroVault associated with this study (https://identifiers.org/neurovault.collection:10447) for relevant maps), but here we highlight one result of interest from an analysis with the vmPFC as a seed region. This analysis interestingly revealed reduced vmPFC-hippocampus (HPC; Right HPC: x, y, z = 22, −14, −20.5; Left HPC: −28, −8, −24) connectivity in older relative to younger adults during reciprocate relative to defect outcomes (see Figure 5 and Table 1).

**Figure 5:**
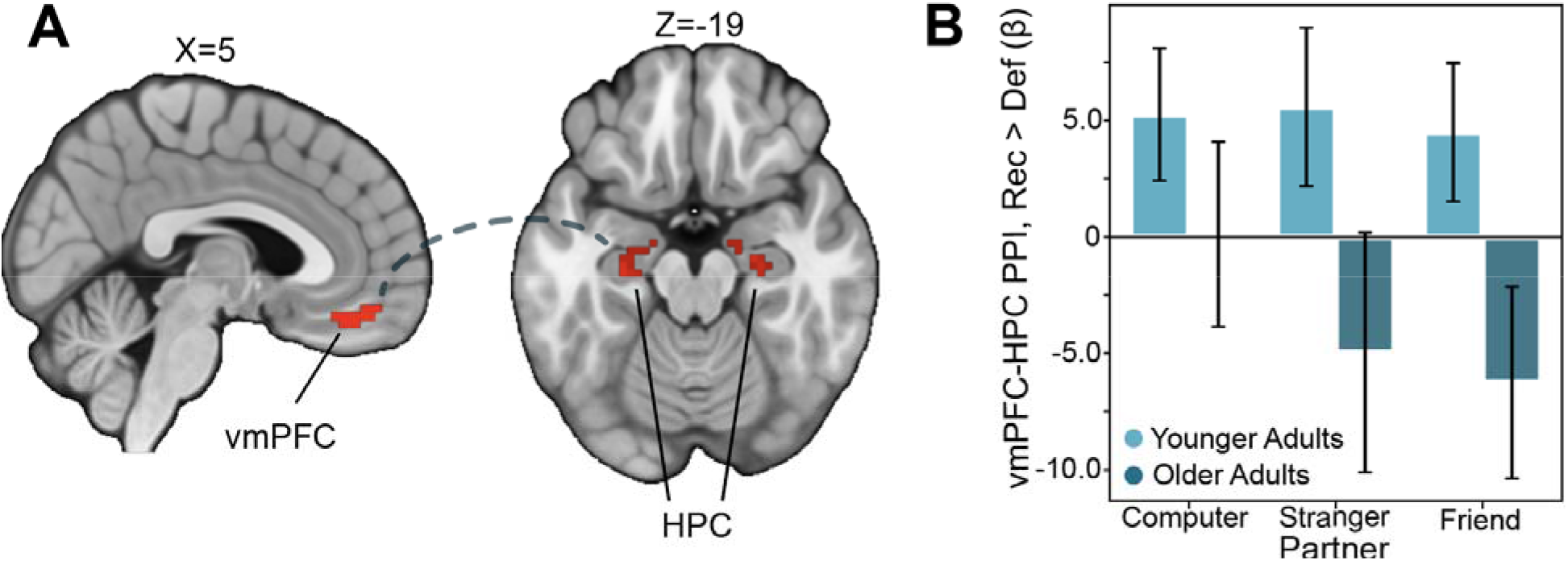
Reduced vmPFC-Hippocampal Connectivity in Older Adults. (A) In an exploratory seed-based PPI analysis, we examined age-related differences in connectivity with vmPFC. We found that older adults exhibited reduced vmPFC connectivity with a hippocampal (HPC) cluster extending into the amygdala during reciprocate relative to defect outcomes (Thresholded: https://identifiers.org/neurovault.image:512046; Unthresholded: https://identifiers.org/neurovault.image:512074). (B) Interrogation of this hippocampal cluster revealed that the pattern was stable across all partner types. We note that Z statistic images were thresholded parametrically (Gaussian Random Field Theory) using clusters determined by Z>3.1 and a (corrected) cluster significance threshold of P=0.05.

## Discussion

Older adulthood is often associated with a heightened focus on experiences with close others in conjunction with shrinking social networks, particularly when emotional goals are prioritized (Carstensen, 1992; Isaacowitz et al., 2021). This changing social world thus colors many daily decisions, including trusting a range of individuals with assets, potentially creating opportunities for exploitation and abuse (Spreng et al., 2016). Although decision neuroscience has made significant progress in characterizing how age-related differences in affect and motivation contribute to decision making (Samanez-Larkin and Knutson, 2015), relatively less is known about age-related differences in social decision-making and social outcome processing. Using an iterated trust game in which participants played with three partners who varied in social closeness with the participant (computer, stranger, friend), we found partial support for preregistered hypothesis 1.1: participants overall invested more with close others, consistent with prior work (Fareri et al., 2015; Webb et al., 2016). Contrary to our predictions, however, we did not observe blunted effects of partner on trust behavior in older relative to younger adults. and in line with pre-registered hypothesis 1.1, younger and older adults demonstrated similar rates of trust with friends and strangers. Interestingly, neural processing of reciprocity and defection varied with respect to age and social closeness. We found partial support for pre-registered hypothesis 2.1, in that striatal responses to reciprocated trust from friends relative to strangers and computers were blunted in older adults relative to younger adults. However, while we did not observe this pattern to be tied to DMN-striatal connectivity (pre-registered hypothesis 2.1), older adults did show enhanced DMN connectivity with the rTPJ during reciprocated trust from friends compared to computers. Taken together, these results suggest that older adults demonstrate altered representations of social closeness within financial exchanges involving trust, which may have downstream, long-term effects on their ability to adapt behavior in social interactions.

Altered recruitment of corticostriatal circuits in older adults when processing reciprocity and defection is partially consistent with earlier work looking at social outcome processing in aging, though we note inconsistencies in the paradigms implemented and patterns of results. Early investigations noted that relative to younger adults, older adults demonstrated altered recruitment of corticostriatal circuits (e.g., dorsolateral PFC, insula) when faced with unfair offers in an ultimatum game relative to younger adults (Harlé and Sanfey, 2012), and an enhanced anticipatory response in the ventral striatum to social rewards in older relative to younger adults (Rademacher et al., 2014). Findings have also shown reduced striatal activation in older adults when exposed to social outcomes that are inconsistent with initial impressions of others (Suzuki et al., 2019). Here, the blunted striatal response to reciprocity from friends relative to strangers and the enhanced insula response to reciprocity from strangers in older adults may suggest an inappropriate weighting of social outcomes that could possibly have implications for the increased susceptibility to financial exploitation in this population, in line with some previous work (cf. Suzuki, 2018). However, we note that the evidence supporting such an interpretation is scant, and requires future testing. An alternative interpretation of these findings might be that positive interactions with strangers may in fact be advantageous. Evidence indicates that the number of weak ties an individual has in older adulthood is inversely related to depressive symptoms and positively associated with positive affect (Huxhold et al., 2020). In addition, recent theories posit that older adults who view aging in a more positive light may see interactions with new acquaintances as opening the door for future engagement and relationships (Huxhold et al., 2022). Thus, enhanced neural responses to positive relative to negative outcomes experienced with strangers in the striatum and insula may reflect an enhanced value placed on interacting with new people.

We do acknowledge that our behavioral results may have limited implications for understanding increased risk for financial exploitation in older adults; however, our imaging results do build on evidence (Spreng et al., 2017; Hughes et al., 2020; Weissberger et al., 2020) suggesting that those older adults who are at heightened risk for financial exploitation show altered activation and connectivity in regions supporting social cognition. For example, decreased mPFC activation and weaker DMN connectivity has been related to diminished mentalizing about unknown others in older relative to younger adults (Hughes et al., 2019; Cassidy et al., 2021). Here, we report enhanced DMN-TPJ connectivity in older adults when experiencing reciprocity from friends relative to computers and strangers. An exploratory analysis revealed that the degree to which older adults invested more with friends over computers was associated with less connectivity during reciprocity (relative to defections) from friends relative to computers; younger adults showed the opposite pattern. Coupled with reported stronger ingroup (vs. outgroup) trust bias in older adults (Cassidy et al., 2020), the tighter integration of TPJ with the DMN during experienced reciprocity with friends here may suggest an augmented ability to incorporate social outcomes into models of known close others. We also found reduced vmPFC-HPC connectivity in older adults during experiences of reciprocity, which, when considered in light of evidence of reduced hippocampal involvement in feedback-based learning and memory (Lighthall et al., 2018) and diminished memory for transgressions in social interactions (Suzuki, 2018) in older adults, may suggest significant changes in the ways in which social and value-related information during trust-based interactions are integrated to inform social learning in aging samples. Future work may examine how age-related connectivity changes during social outcome processing may relate to neural differences during trust-related decision-making as a function of social closeness.

We note that our results merit further consideration of a few important caveats. First, we observed no age-related differences in willingness to trust others as a function of social closeness, which stands in contrast to the patterns observed in the imaging results. This could be partly attributed to our small sample size (we were powered to detect moderate to large effects). It is also possible that age-related differences in behavior are more subtle, potentially requiring advanced computational methods to estimate latent factors in the decision-making process (Miletic et al., 2020), or additional clinical assessments of cognitive and neural decline (e.g., Diffusion Tensor Imaging) An alternative possibility is that because the choices in this task were designed to assess levels of trust in one given individual at a time, there were benefits to placing trust in a stranger, so as to learn about them (Isaacowitz et al., 2021). While such an interpretation is at odds with some theories of social reorientation in older adulthood (Carstensen, 1992), it is in line with suggestions of the benefits of maintaining less close relationships in older adulthood (Huxhold et al., 2020), which may serve contextual goals that differ across individuals (Huxhold et al., 2022).

We also note that inconsistencies are often reported with respect to age-related differences in trust decisions—older adults have been observed to both invest more with trustees (Bailey et al., 2016), and to demonstrate no differences relative to younger adults in investment behavior (Bailey et al., 2015)—while other findings note that older adults may be less able to effectively adapt behavior after negative (i.e., betrayal of trust) social outcomes (Bailey et al., 2019; Frazier et al., 2021). These differences may be in part due to variability in experimental design. Future work may implement alternative two-stage designs (Daw et al., 2011) in which older adults have the option to choose to interact on a given trial with a close or unknown other before deciding how much to invest, which may help disentangle some of these inconsistencies. In addition, the effects of social closeness on behavior could be influenced by other aspects of the relationships that we did not assess, including social network size (Kwak et al., 2018) and relationship quality (Santini et al., 2015), or by other aspects of cognition (i.e., episodic memory; see Suzuki, 2018). Additionally, all of our RT-related analyses were exploratory and not tied to our primary research questions. Future work will have to consider the effects of RT more thoroughly.

Last, we note a few considerations regarding the observed age-related differences in brain activation and connectivity. We attempted to address known potential confounds by including covariates such as gender, head motion, data quality, and response time in analyses of age-related differences. Still, our results could be linked to cohort effects specific to the recruited groups of older and younger adults, which should be addressed in future studies with larger sample sizes that track changes across time. In addition, it is possible that other variables, such as white matter integrity and vascular health (Prins & Scheltens, 2015), may have contributed to the age-related neural differences we observed. Indeed, changes in white matter tracts assessed via Diffusion Tensor/Weighted Imaging have been associated with later diagnosis of neurodegenerative disorders associated with cognitive decline (e.g., Alzheimer’s Disease) (Benear et al., 2020); future studies could assess these types of data to identify potential neuro-predictive markers associated with future risk for exploitation in social settings. Evidence also points to dynamic age-related changes in structural and functional connectivity that may underlie aspects of cognitive decline (Madden et al., 2017). Though we were unable to assess these variables in our current study, it is certainly possible that they may be driving the age-related neural differences reported here. Relatedly, while encouraging, we do note that a number of our imaging findings were reduced in strength with the addition of covariates into our analyses. We also acknowledge that the finding of enhanced DMN-TPJ connectivity emerged from a whole-brain contrast of outcomes with friends relative to the computer, not friend relative to stranger. While we did not observe significant whole brain connectivity results for the latter contrast, we believe that the friend > computer analysis is still informative as it highlights a clear social relative to non-social difference. We also did not collect information regarding race, or socioeconomic status, which could be additional factors driving or contributing to age-related neural differences in our sample. Future work should take these additional factors into account, and aim to obtain larger, longitudinal samples to specifically interrogate trajectories of age-related changes in trust behavior and its neural correlates.

Despite these limitations, our results support the conclusion that older adults exhibit altered neural responses to social closeness during trust-based social interactions. While our study represents only an early step at characterizing age-related differences in trust and reciprocity, these results lay the groundwork for future studies to unpack implications for vulnerability to social victimization and financial exploitation among the elderly (Lichtenberg et al., 2013; Lichtenberg, 2016; Spreng et al., 2016). Although financial exploitation is a multifaceted issue that can occur in a wide range of scenarios (Beals et al., 2015; Nguyen et al., 2021), many instances of financial exploitation may stem from placing too much trust in others (Shao et al., 2019; Nguyen et al., 2021). We speculate that future efforts to understand the moderating effect of social closeness on decision making and reward processing in older adults may ultimately better inform novel risk factors for financial exploitation.

## Acknowledgments

The authors thank Elizabeth Beard for assistance with task coding, Dennis Desalme, Ben Muzekari, Isaac Levy, Gemma Goldstein, and Srikar Katta for assistance with participant recruitment and data collection, and Jeffrey Dennison for assistance with data processing. DVS was a Research Fellow of the Public Policy Lab at Temple University during the preparation of this manuscript (2019-2020 academic year).

