## Supplementary Information for "Age-Related Differences in Ventral Striatal and Default Mode Network Function During Reciprocated Trust" (from another version of this preprint: version 3, 2022-03-02)

### Supplemental Methods

#### Group-Level Independent Component Analysis

We used Probabilistic Independent Component Analysis (Beckmann & Smith, 2004) as implemented in MELODIC (Multivariate Exploratory Linear Decomposition into Independent Components) Version 3.15, part of FSL (FMRIB's Software Library, [www.fmrib.ox.ac.uk/fsl](http://www.fmrib.ox.ac.uk/fsl)) to identify networks of co-activation within our data. As part of the MELODIC preprocessing, the following procedures were applied to the input data: masking of non-brain voxels; voxel-wise de-meaning of the data; normalisation of the voxel-wise variance. Next, pre-processed data were whitened and projected into a 114-dimensional subspace using probabilistic Principal Component Analysis where the number of dimensions was estimated using the Laplace approximation to the Bayesian evidence of the model order (Beckmann & Smith, 2004; Minka, 2000). The whitened observations were decomposed into sets of vectors which describe signal variation across the temporal domain (time-courses), the session/subject domain and across the spatial domain (maps) by optimising for non-Gaussian spatial source distributions using a fixed-point iteration technique (Hyvarinen, 1999). Estimated Component maps were divided by the standard deviation of the residual noise and thresholded by fitting a mixture model to the histogram of intensity values (Beckmann & Smith, 2004).

To facilitate comparison with our primary network connectivity results, we used a spatial correlation analysis to systematically assess the similarity between each of the 114 spatial maps from MELODIC and each of the 10 canonical maps from prior work (Smith et al., 2009). See Supplementary Table 1.

#### Deviations from Pre-registration

*We note that our analyses deviated slightly from those defined in our pre-registration in the following ways. To test the hypothesis that older adults would demonstrate reduced levels of trust relative to younger adults (H1.1), we conducted a repeated measures ANOVA instead of a mixed-effects regression. We acknowledge an error in this pre-registration––the mixed effects regression examining effects of offer type on ‘rejection rates’ was accidentally copied from a separate pre-registration from our group examining ultimatum game behavior (INSERT LINK). The present study did not examine varying initial offer types presented to participants, nor was the dependent variable in this task rates of rejection, but rather investment behavior. As per our previous work examining investment behavior in a trust game (Fareri et al., 2012; Fareri et al., 2015), we thus decided to conduct a repeated measures ANOVA.* In order to examine the relation between self-reported relationship closeness and reward-related neural responses (H2.1), we also included social closeness as a covariate in an offline analysis with extracted parameter estimates, as opposed to at the whole brain level. This was done in order to maximize statistical power and preserve degrees of freedom at the whole-brain level. Given that we did not observe evidence for age-related DMN-striatal connectivity (H2.2), we did not test H2

Additionally, we conducted a number of exploratory analyses based on both previous literature as well as patterns that emerged in the data which complement our pre-registered questions. We conducted a task-based generalized network PPI analysis to examine connectivity of the Executive Control Network (ECN) during the processing of trust game outcomes, based on our previous work (Fareri et al., 2020; Utevsky et al., 2017). Last, we conducted whole-brain seed-based PPI analyses using the fusiform face area (FFA), ventromedial prefrontal cortex (vmPFC) and ventral striatum, TPJ and PCC as seed regions in order to investigate age-related differences in responses to trust game outcomes as a function of social closeness.

### Supplementary Results

We extracted connectivity estimates between the DMN and TPJ cluster that emerged in our whole brain analysis reported in the main manuscript and conducted a 2 (age group) x 2 (partner: computer, friend) repeated measures ANCOVA including tsnr and mean framewise displacement as covariates. Results of this control analysis paralleled the results reported in the main manuscript: a significant age group x partner interaction emerged in DMN-TPJ connectivity (*F_(1,44)_* = 9.10, *p* = .004) such that older adults show enhanced connectivity during reciprocate relative to defect outcomes from friends compared to computers, whereas younger adults demonstrate the opposite effect. We also extracted connectivity estimates between the ECN and an insula cluster that emerged in our whole brain analysis reported in the main manuscript and entered them into a 2 (age group) x 3 (partner: stranger, friend) repeated measures ANCOVA with tsnr and mean framewise displacement as covariates. This analysis revealed a trending partner x age group interaction (*F_(1,44)_* = 3.03, *p* = .089), such that older adults, compared to younger adults, show enhanced connectivity during defect relative to reciprocate outcomes from friends relative to strangers. These results parallel those reported in our main ECN connectivity analysis, though we again note that this supplemental finding is a trend.

In an exploratory analysis, we examined links between network connectivity and trust behavior. As detailed in the main text, we found that age moderated the relation between DMN-TPJ connectivity and trusting behavior across friend and computer partners (Supplemental Figure 1).

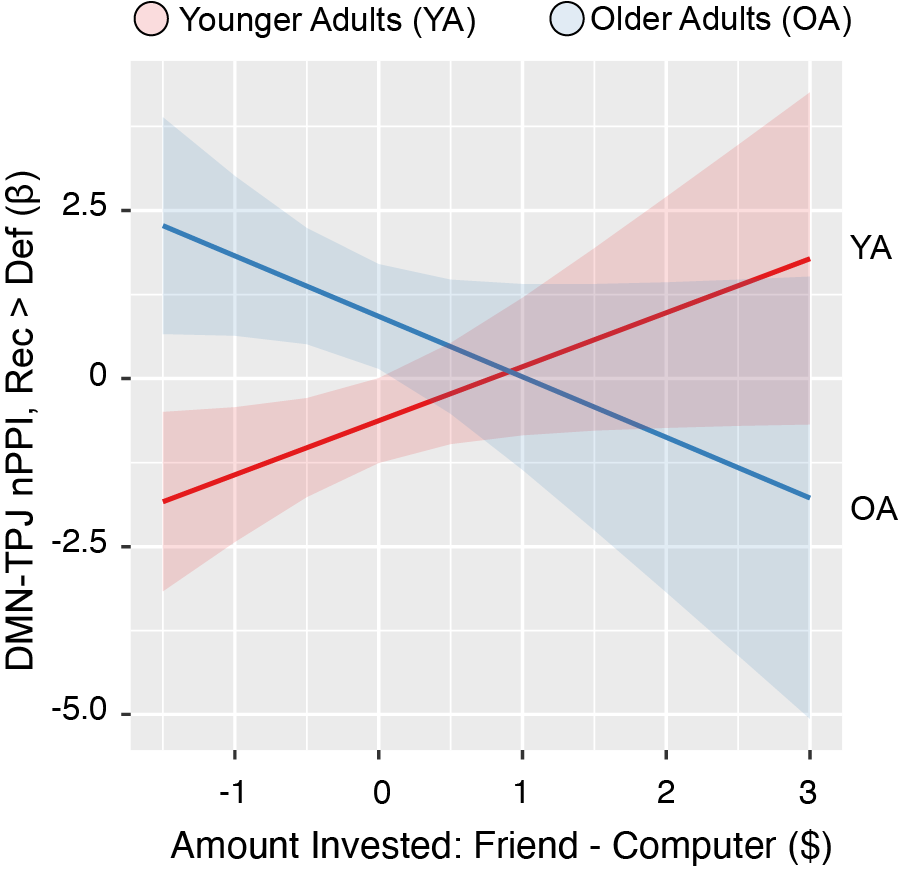

**Supplemental Figure 1: Moderating effects of age on brain-behavior relations.**

### Supplementary Tables

#### Supplementary Table 1

Linking MELODIC output and canonical network maps in prior work (Smith et al., 2009). The “melodic_IC” row indicates the independent component number from our analysis. The second row indicates the correlations between each of the canonical network maps indicated in the columns (note: Smith09_04 is DMN and Smith09_08 is ECN). To illustrate the selectivity of these matches, the next four rows represent the mean, standard deviation (std), minimum (min), and maximum (max) correlation values across other components.

|  | **smith09_01** | **smith09_02** | **smith09_03** | **smith09_04** | **smith09_05** | **smith09_06** | **smith09_07** | **smith09_08** | **smith09_09** | **smith09_10** |
| --- | --- | --- | --- | --- | --- | --- | --- | --- | --- | --- |
| **melodic_IC** | 27 | 31 | 31 | 48 | 50 | 13 | 43 | 16 | 19 | 26 |
| **corr with Smith09** | 0.68 | 0.46 | 0.47 | 0.70 | 0.41 | 0.38 | 0.38 | 0.45 | 0.52 | 0.54 |
| **mean corr (other)** | 0.01 | 0.00 | 0.01 | 0.00 | 0.00 | 0.01 | 0.00 | 0.01 | 0.01 | 0.00 |
| **std corr (other)** | 0.05 | 0.05 | 0.06 | 0.05 | 0.06 | 0.07 | 0.07 | 0.07 | 0.06 | 0.06 |
| **min corr (other)** | -0.11 | -0.13 | -0.14 | -0.18 | -0.12 | -0.09 | -0.18 | -0.09 | -0.10 | -0.15 |
| **max corr (other)** | 0.33 | 0.44 | 0.26 | 0.18 | 0.28 | 0.37 | 0.35 | 0.32 | 0.28 | 0.29 |
